# Prey refuge and morphological defense mechanisms as nonlinear triggers in an intraguild predation food web

**DOI:** 10.1101/2020.02.04.934588

**Authors:** J. P. Mendonça, Iram Gleria, M. L. Lyra

## Abstract

Intraguild predation (IGP) is a type of interaction in which a top predator simultaneously competes and predates an intermediate prey that shares a third prey species with the top predator. While common in nature, most theoretical population dynamics models proposed in the literature predict that this three species interaction usually leads to extinction of the intermediate prey population. Predator-induced defense as well as refuge mechanisms are widely seen in these systems and should be incorporated in IGP models to promote coexistence. With this aim, we introduce a nonlinear response to the predation of IG-predator on IG-prey modelling both prey refuge and morphological defenses. The phase diagram of species coexistence is obtained as function of the attack efficiency and the degree of nonlinearity of the defense mechanisms. Further we show how the nonlinearity affects the equilibrium populations. We unveil that there is an optimal nonlinearity at which the convergence towards the stationary coexistence regime is the fastest.

## 1. Introduction

Intraguild predation (IGP) is a class of omnivorous food web that combines predation and competition. Two species exploit the same environmental resource and one of them (the IG-predator) predates the other (IG-prey) [1]. Such kinds of food webs are theoretically predicted to decrease the potential of being stable, thus leading to extinction of both species [2]. Notwithstanding, intraguild predation is widespread [3, 4, 5, 6] and this ubiquitous interaction is commonly observed in nature. There are several biological mechanisms to promote the persistence of the food web, namely defenses induced by the presence of the IG-predator. Experimental and observational studies showed different dynamical patterns in food webs where inducible defenses are present [7]. Such defenses bring into play changes in morphology [8, 9, 10, 11], life-history [12] and prey behavior [13, 14], to name a few.

Among the systems in which the predator-prey interactions allows for the growth of the intermediate prey, we highlight those on which a morphological defense is induced by the predator presence. In a recent paper, Kratina et al. [7] studied the dynamical properties of a food web consisting of the resource food, a type of unicellular algae, *Rhodomonas minuta*, which is consumed by both the turbellarian flatworm, *Stenostomum virginianum*, and the hypotrich ciliates of genes *Euplotes* spp, while the last is also consumed by the flatworm. In the presence of a large predators density, the Euplotes changes its morphology increasing substantially the body size, avoiding predation by the flatworm which is a gape-limited predator. Consequently, in the defensive form, the Euplotes are too big to gobble and could promote the species survival and then the persistence of the food web.

Preys can also use a refuge strategy to avoid predation. Whitlow et al. [15] studied a system of soft-shell clams, *Mya arenaria*, in response to introduced green crab predators, *Carcinus maenas*. Soft-shell clams increases pressurization inside the mantle cavity following closure of the siphon and contraction of the valves [16]. Using this burrowing mechanism, clams eject water through its pedal gape while using the foot to grip the sediment. Foraging crabs spend time searching and excavating clams, so that clams could be less vulnerable burrowing deeper in the sediment [17]. As excavation can build a refuge from predation, these results show that this mechanism is an inducible defense against predatory crabs. Simplification on the IGP model to a two-species community with an unlimited resource food proved too simple on account of the low complexity of the system [2, 18, 19]. However, when it is required to take into account the temporal evolution of the resource food, the proper modelling of inducible defenses has been a challenging task [20, 21, 22, 23, 24, 25].

Dealing with two species only, Vos et al. [26] showed, with realistic parameters, how consumer-induced defenses are important to bottom-up control using food chain models. Ramos-Jiliberto et al. [27] explored pre- and post-encounter inducible defenses in predator-prey systems that have similar dynamical effects and showed that inducible defenses prevent the paradox of enrichment and should increase the stability. Using dynamic-optimization models, Clark and Harvell [28] studied the cost of defense. Addition of a “delayed” term in reproduction was shown to favor a higher allocation to defense. A model for the predator-induced defense that analyzes the cost-benefit of the defensive morph was introduced in *D. pulex* by modifying the life-time of both induced and normal morphologies for different predation rates of *Chaoborus*, showing that the inducible defense could be less advantageous in certain cases [29]. Those studies are applied to the complexity of populations dynamics models with up to two species only or with a weak interaction with the resource food [2, 18, 19, 30].

When increasing the complexity of the food webs, with strongly interacting species, most models become more and more unstable. Stability requires that the IG-prey be a better competitor than the IG-predator [31, 32]. In order to include inducible defenses to obtain the species coexistence in an IGP model, Nakazawa et al. [21] incorporated predator-specific defense adaptation of the resource food against both IG-prey and predator, showing clearly that this mechanism favors coexistence. A key study made by Holt and Huxel [22], focused on the three-species community. The authors introduced other types of community modules as alternate prey. They brought to light the great influence of alternate prey in the persistence of the food web as well as presented a discussion concerning the models with Lotka-Volterra-like functional responses, which are clearly limited. The authors emphasized that it should be necessary to use other nonlinear response models [33, 34] to properly deal with the defense mechanism. Regarding the nonlinear functional responses to model these interactions, there are several proposals using distinct realistic functions with different interpretations such as Holling Type II [35], Type III [36], Beddington–DeAngelis [37, 38] and ratio-dependent [39, 40, 41] functionals. Drossel et al. [41] showed via evolutionary models, that ratio-dependent nonlinear functional responses enables the emergence of large and complex food webs with stable structures.

In this work, we introduce two kinds of inducible defenses in an IGP model, refuge and morphological, treating the dynamical evolution of the three-species community. We insert nonlinear triggers such as prey-dependent and predator-dependent functional responses, showing that it is possible to switch from extinction of the IG-prey (without defenses) to the persistence of the food web (within defenses). We also build phase diagrams in parameter space spanned by the degree of nonlinearity associated to the defense mechanism and attack efficiency of the IG-predator to the IG-prey. By determining how the rate of convergence to the coexistence regime depends on the degree of nonlinearity introduced into the IG-prey/IG-predator interaction we summarize all long-term regimes. Finally, modelling the refuge mechanism, we show the effect of a saturable nonlinearity on the phase diagram.

### 2. IG predator-prey model

The basic interaction of the intraguild predation system considers a resource population *R* that is consumed by both an herbivorous population *E*, the IG-prey, and an omnivorous population *S*, the IG-predator. The latter also consumes the IG-prey, see Fig. 1. The resource food population *R* increases by itself via photosynthesis and nutrients such as nitrogen, phosphorus, and potassium. Since the nutrients and sunlight areas are limited sources of food, they compete with each other and for that reason, we assume that it is increasing logistically. For the intermediate/top species, we can assume that the populations find plentiful food at all times and the food supply depends only on the size of the prey population. Also, the rate of population changes is proportional to its size. The above assumptions just describe a system that could be entirely described by Lotka-Volterra interactions. In nature, however, many food webs persist towards more complex interactions such as anti-predator mechanisms which are our basic interest in this study. In what follows, we take into account the IG-prey defense reaction based in biologically motivated mechanisms.

**Figure 1:**
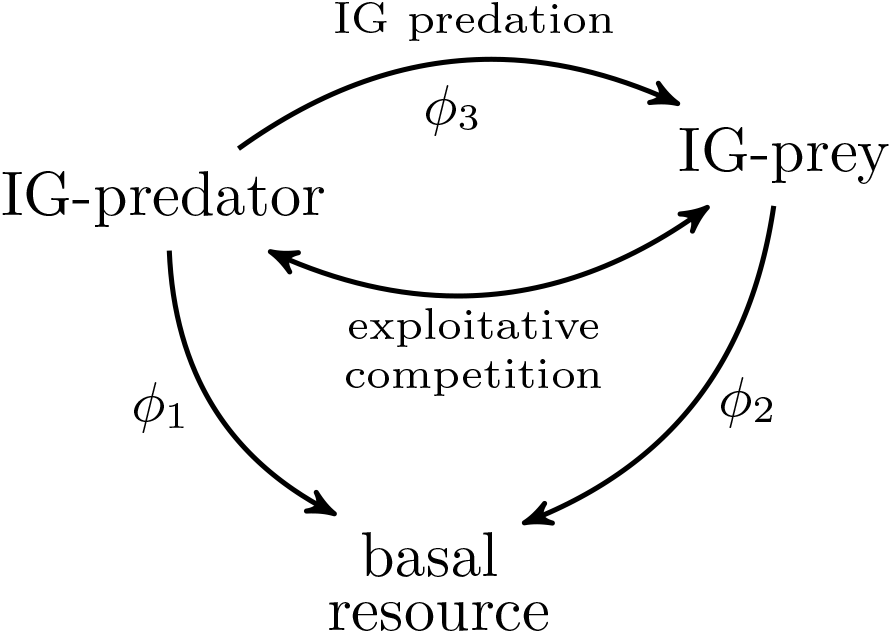
Food web scheme. Arrows indicate that one species (head arrow) has a detrimental effect caused by a second one. Predations are balanced by attack efficiencies *ϕ*.

The differential equations for the above described model are:

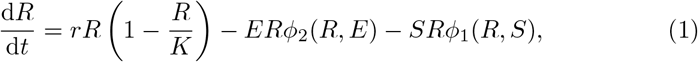

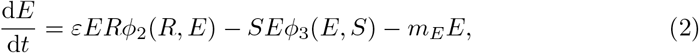

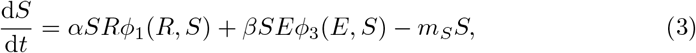

where *ϕ_i_* = *a_i_* (*i* = 1, 2) are Lotka-Volterra-like attack efficiencies and *ϕ*_3_ accounts for the intraguild attack efficiency. We incorporate anti-predator mechanisms by assuming it to act as a nonlinear trigger. In what follows we will model two distinct defense mechanisms, namely the refuge hiding of small surviving populations of the IG-prey and the morphological defense reaction of the IG-prey population triggered by large populations of IG-predators.

### 2.1. Refuge mechanism

Prey species that change behavior in response to a predator finding or building places to refuge can be modelled by a nonlinear prey-dependent response. Often one assumes a certain number of hiding places or some potential places to burrow deeper. A greater proportion of the prey can find these places as prey number becomes smaller. According to this reasoning, the attack efficiency shall decrease when the IG-prey population approaches extinction. This feature can be effectively incorporated in the attack efficiency *ϕ*_3_ by assuming it to become vanishingly small as *E →* 0. Here, we model such refuge mechanism of the IG-prey by a nonlinear trigger assuming *ϕ*_3_ to have the form

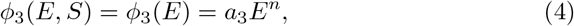

where *n* is the degree of nonlinearity of the IG-predator/IG-prey interaction.

Behavioral responses to predators faces a trade-off between predation and feeding. In the case of soft-shell clam, its feeding is not affected if its siphons are long enough to get to the surface [15]. Our assumptions are plausible for soft-shell clams, *Mya arenaria*, and differs from other species where their larger body size is a refuge from predator. When the nonlinearity becomes increasingly stronger, the consumption of IG-prey rapidly approaches zero when the IG-prey population *E* is low. Nevertheless, the nonlinear response turns the IG-predator attack more efficient for large densities of *E*. In fact, assuming that the IG-prey has only this anti-predator mechanism, the attack efficiency of the predator must be increased by the abundant presence of the prey, since they are vulnerable when in great quantity. Note that, larger values of *n* leads to a stronger suppression of the attack efficiency for low IG-prey densities, as shown in Fig. 2. For large *E* populations, this nonlinear coupling actually favors the consumption of *E*. This points to a relation between cost and benefit, that plays a key role in the dynamics and is a fundamental element to reach the persistence of the food web. However, as the population of *E* is limited by the availability of resources, the regime with very large *E* population is not reached. Therefore, there is no point where the IG-predator will not be interested in consuming the basic resource *R* due to an excessive abundance of IG-prey.

**Figure 2:**
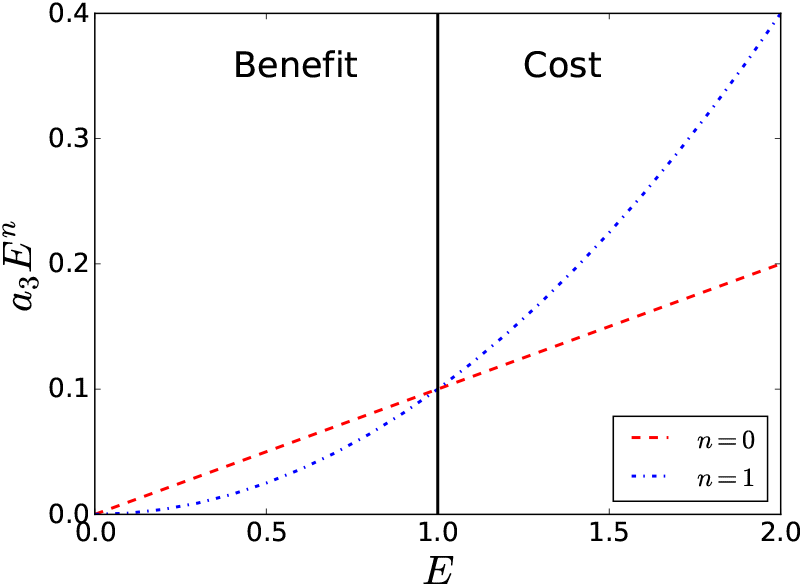
Illustrative plot of the linear and nonlinear IG-prey/IG-predator interaction with anti-predator induced refuge. Notice that the nonlinearity strongly suppresses the attack efficiency at low *E* populations. Parameters: *a*_3_ = 0.1.

### 2.2. Morphological defense mechanism

The presence of a large density of IG-predators creates chemical cues that cause the IG-preys to invest energy into defense [42, 43]. In morphology-induced defense reaction, the IG-prey can increase in size preventing its consumption. The change in body size brings out negative influences to the predator. As pointed out by Kratina et al. [7], when *S* is large the defense becomes more effective and can promote the persistence of the food web. The defensive mechanism can be incorporated in the attack efficiency by assuming it to become vanishingly small at very large IG-predator populations. Here, we can also model this morphology-induced defense reaction as a nonlinear trigger by assuming *ϕ*_3_ to have the form

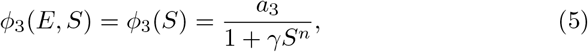

where *n* is the degree of nonlinearity of the IG-predator/IG-prey interaction which has a saturation strength *γ*. Therefore, for large predator densities with *γS^n^ »* 1 the defense mechanism is activated and the attack efficiency becomes vanishingly small. We are not introducing costs to employ the inducible defense. However, it is known that inducible defenses display several biological trade-offs such as fitness, feeding rate and metabolic costs, to name a few. The main assumption is that the predation risk cannot be high enough and, as theorized by Ramos-Jiliberto et al. [44], with variable resources, inducible defenses produce small costs when predation risk is lower.

Fig. 3 shows the functional response for different values of nonlinearity degree. With a high predator abundance the attack fades away. We see that the attack quickly vanishes with increasingly values of *n*. The linear functional response is recovered for *γS^n^ «* 1.

**Figure 3:**
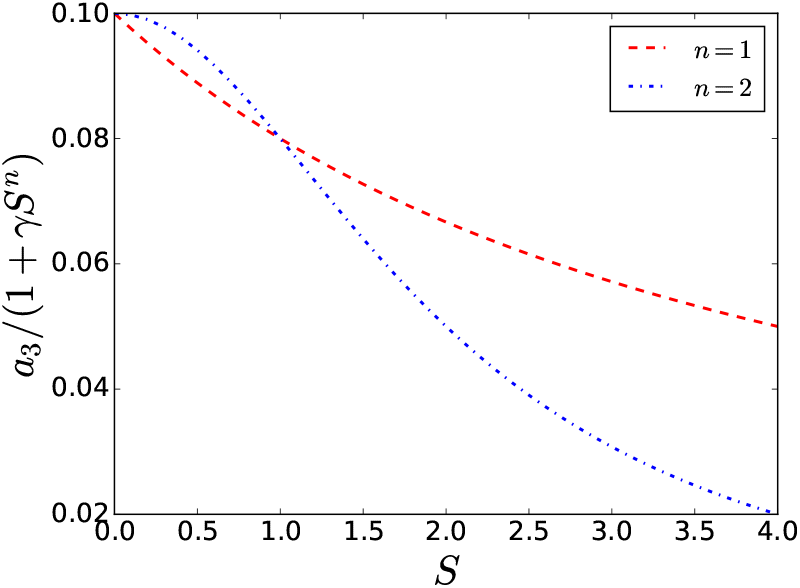
Illustrative plot of the nonlinear IG-prey/IG-predator interaction with morphological inducible defense. The attack efficiency is lower for higher IG-predator population abundance. Parameters: *a*_3_ = 0.1 and *γ* = 0.25.

## 3. Results

### 3.1. Persistence of the food web

An analysis of the equilibrium points of the above dynamical system shows that the nonlinear defense has the potential to induce a stability switch of the boundary fixed point (*R*,* 0*, S\**). Solving **F**^*\**^ = 0, **F** being the three-component vector of differential equations of the model presented and **F**^*\**^ its value evaluated in the fixed point, we obtain the equilibrium points of the present dynamical system. The stability is determined by the eigenvalues *ω* of the Jacobian matrix *J* (*R*, E*, S\**) by det*|J − ω**1**|* = 0, where ****1**** is the identity matrix. The stability is determined by the real part of the eigenvalues, say Re*{ω}*. Explicitly we have:

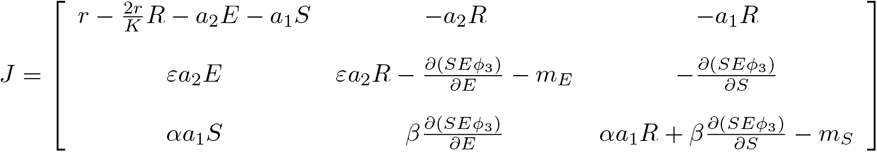

First, we analyse the stability of the fixed points for the refuge model. There are two different Jacobians for *n* = 0 and *n* ≠ 0 when *E\** = 0. For the boundary fixed point 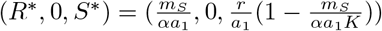 we obtain:

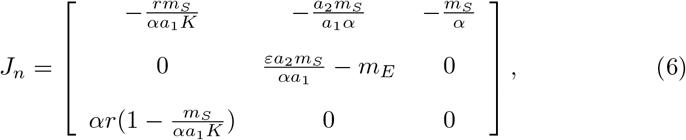

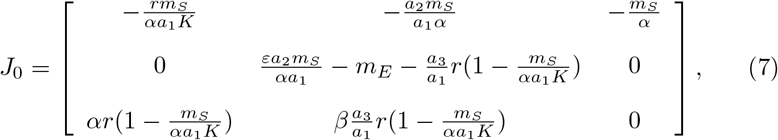

with the eigenvalues given by:

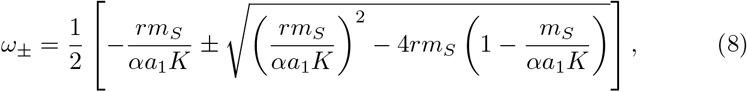

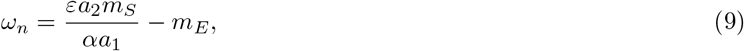

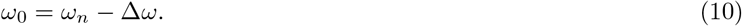

Here 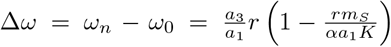 bring us the difference in stability between the linear and nonlinear response models. Δ*ω* will tell us if it is possible to have a stability switch by introducing the nonlinear inducible defense on the form of Eq. 5. In fact, the fixed point is stable if Re*{ω} <* 0 for all *ω* and unstable otherwise.

Some of the above features related to the stability of the fixed points change when the attack efficiency is compromised by a morphological defense response. In this case, the stability of (*R*,* 0*, S\**) is determined by the *n*-independent eigenvalues *ω_±_* and

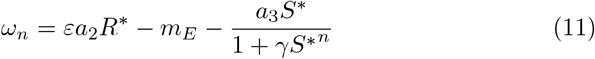

where 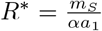 and 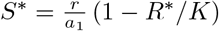.

The anti-predator mechanism cannot affect the food web when the IG-predator is excluded. Therefore this point feels no change and it is equal to the IGP Lotka-Volterra model, which is a reasonable result regarding the fact that an anti-predator mechanism affect the food web only when the predator is present.

The following numerical analysis was made using the set of parameters shown in Table 1. We find that the Re*{ω_±_} <* 0, *ω_n_ >* 0 and Δ*ω > ω_n_*. Thus in the basic linear model (*n* = 0) the boundary point (*R*,* 0*, S\**) is stable. With the nonlinear trigger we get similar outcomes for both mechanisms. The fixed point with *E* excluded becomes unstable within certain values of *n*. One reasonable assumption is that the stability switch of the boundary fixed point is actually followed by stability switches of other (possibly internal) fixed points. In Fig. 4 we show via numerical solutions of the dynamical equations that the stable point (*R*,* 0*, S\**) becomes unstable (or infeasible) whereas the unstable (or infeasible) point (*R*, E*, S\**) switches to stable when we introduce the nonlinear defense. We also show in Fig. 4c-d the possibility of switching from fixed point to a limit cycle just by varying the nonlinearity degree. This feature obtained from the morphology-induced defense model is not presented by the refuge model where the oscillations are rapidly damped. Another interesting fact is that even if the IG-prey is not a better competitor (*a*_2_ *< a*_1_), we still get similar results, reaching the persistence of the food web. We are thus led to conclude that a nonlinear response associated to an anti-predator mechanism is a fundamental tool to preserve the food web.

**Table 1:** Parameters used in the present study. They were chosen to have the boundary fixed point (*R*,* 0*, S\**) stable in the regime of linear IG-prey/IG-predator interaction.

|  | Meaning | Value |
| --- | --- | --- |
| $r$ | Intrinsic growth rate | 1.5 |
| $K$ | Carrying capacity | 5 |
| $\varepsilon, \alpha, \beta$ | Conversion efficiency | 1.2, 1, 0.75 |
| $a_{1,2,3}$ | Attack efficiency | 0.75, 1.5, 0.1 |
| $m_E$ | Death rate of E population | 0.25 |
| $m_S$ | Death rate of S population | 0.15 |
| $\gamma$ | Saturation strength | 0.25 |
| $n$ | Degree of inducible defense | - |

**Figure 4:**
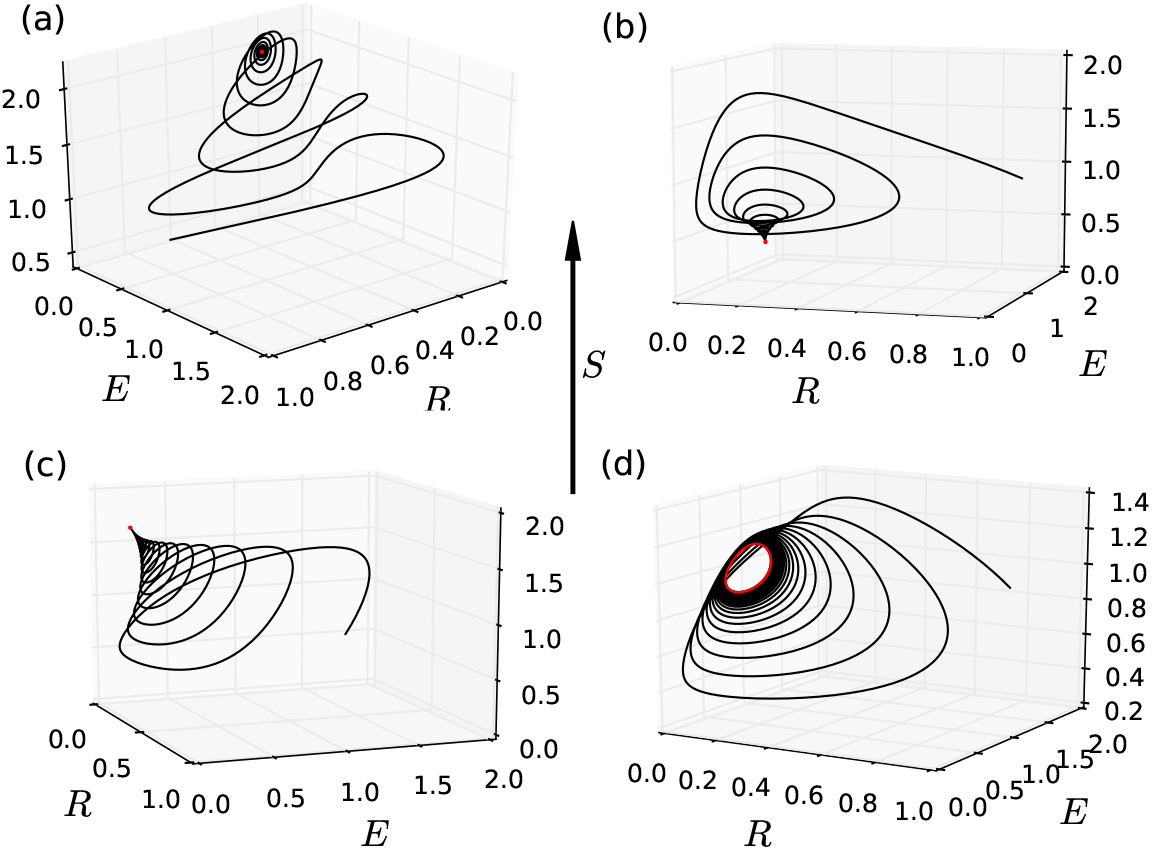
Phase space for (a) Linear case for which we recover the basic model and the *E* population is extinct; (b) Morphological defense with *n* = 2 showing a stable internal fixed point; (c) Refuge defense with *n* = 2 showing a stable internal fixed point; (d) Refuge defense with *n* = 15 showing a limit cycle with persistent oscillations. Parameters are given in Table 1.

The population densities in equilibrium (fixed point) depend on the degree of nonlinear defense besides the other usual dynamical parameters. Fig. 5 shows us the dependence of the stationary populations in the internal fixed point (coexistence point) against the degree of nonlinearity *n*. Note that there is a characteristic value of the degree of nonlinear defense where a stability switch occurs. With a refuge defense mechanism, we see that above this value the equilibrium density of the IG-predator population *S* is zero, the defense is so effective that it can lead to the extinction of the population of the IG-predator. This counter-intuitive result can be understood by recalling that the IG-predator population only persists without attacking the IG-prey if *αR*a*_1_ *> m_S_*. An efficient hiding defense increases the IG-prey population thus leading to a smaller availability of the basic resource which is given by *R\** = *m_E_/∊a*_2_ in the fixed point with *S\** = 0. Therefore, the condition for the extinction of IG-predators when its attack to the IG-prey is inefficient is *αm_E_a*_1_*/∊m_S_a*_2_ *<* 1 that is fulfilled for the model parameters we used. In the case of morphological defense, shown in Fig. 5b, coexistence is developed above a characteristic degree of nonlinearity. There is also a second stability switching point above which the system develops sustainable oscillations over a limit cycle. In this regime, the reported populations are averages over a cycle.

**Figure 5:**
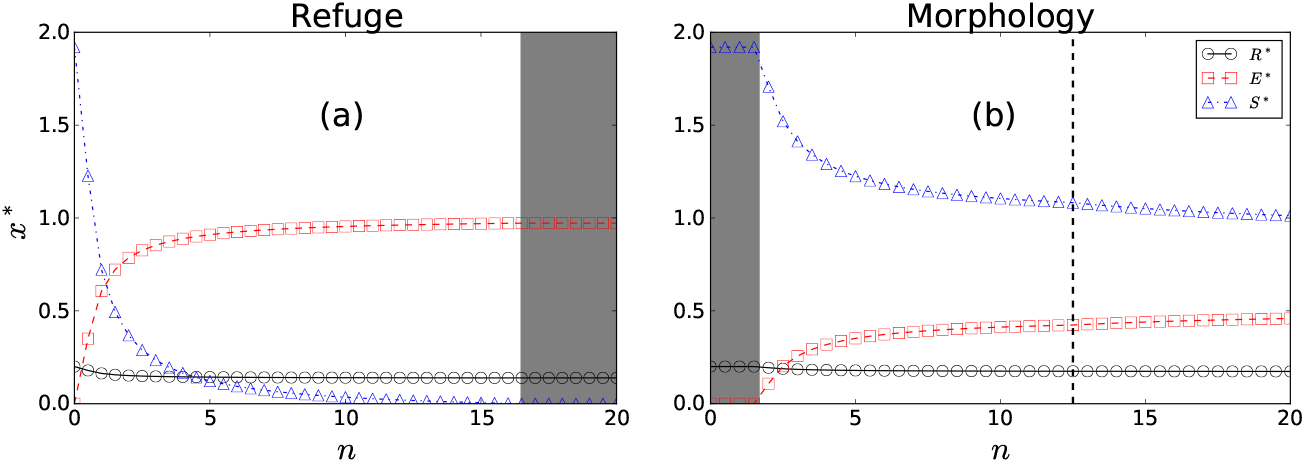
Dependence of the populations at the stable equilibrium point on the degree of nonlinear defense. (a) Refuge defense: Notice that any degree of nonlinearity leads to the persistence of the IG-prey population. The food web is stable for 0 *< n < n\**. Above *n* the IG-predator population becomes extinct. (b) Morphological defense: The food web is stable for any *n > n\**. Below this value the IG-prey population is still extinct. There is a second stability switch above which the system evolves towards a limit cycle. In this regime we report the average populations over a cycle. Here *x* represents either *R*,*E* or *S*. Parameters are given in Table 1.

The characteristic nonlinearity *n\** leading to the stability switch between the coexistence and non coexistence phases was determined analytically by analyzing the linear stability of the *n*-dependent boundary fixed point, which has *S\** = 0 for the refuge defense or *E\** = 0 for the morphological defense as seem above. We determined under which condition its stability changes. For the refuge model, it is given by

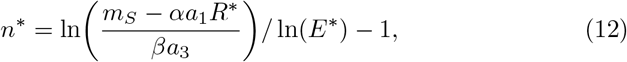

where 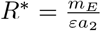 and 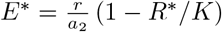 For morphological defense one gets

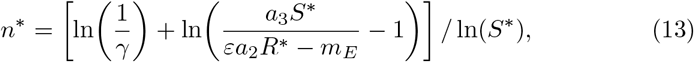

where 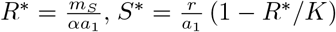 and *γ* ≠ 0. For the set of parameters used in our numerical work, this characteristic nonlinearity for the refuge defense model is *n* ≃* 16.48, with coexistence persisting for 0 *< n < n\**. For the morphological defense model one finds *n* ≃* 1.67 above which all three species coexist.

The attack efficiency of the top predator on the intermediate prey is essentially modulated by two parameters: the linear attack efficiency *a*_3_ and the degree of nonlinear inducible defense *n*. With a fixed *n*, *a*_3_ controls the attack efficiency while, with a fixed *a*_3_, *n* controls the defense efficiency (or the attack inefficiency). Based on the above phenomenology, we built a phase diagram by varying these two parameters such that the *n\** curve separates the regime of coexistence of all species and the extinction of one species (see Fig. 6). However, it is important to stress that, in the case of refuge mechanism, the IG-prey population in equilibrium continuously decreases as *n →* 0 (see Fig. 5a). There is so much refugee prey for *n > n\** that the attack becomes very inefficient. Assuming the IG-predator could not utilize prey-switching or adaptive feeding, this leads to the extinction of the IG-predator population due to a reduced availability of basic resources. For the morphological defense, the IG-prey population becomes extinct for *n ≤ n\** (see Fig. 5b).

**Figure 6:**
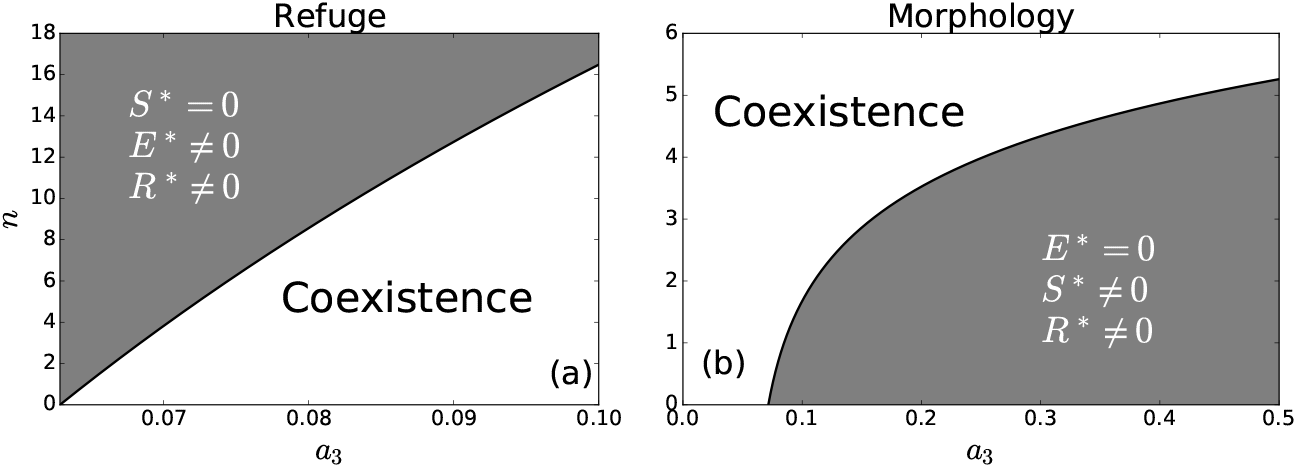
Phase diagram in the parameter space spanned by the linear attack efficiency *a*_3_ and nonlinearity *n*. The solid black line shows the characteristic nonlinearity of the defense *n\** which separates the phases. In the shaded region the food web is broken. (a) Refuge defense: For *n > n\** the IG-predator becomes extinct; (b) Morphological defense: For *n < n\** the IG-prey becomes extinct. Parameters are given in Table 1.

### 3.2. Rate of convergence to the coexistence equilibrium

The dynamical behavior of the system can be written in terms of the eigenvectors of the Jacobian matrix 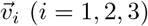. Near the fixed point (in a linear approximation) we have *v_i_*(*t*) *≈ v_i_*(0)*e^ω^it*, where *ω_i_* is the corresponding eigenvalue. In populations dynamics terminology, we say that the eigenvectors are the principal directions to the dynamics of the interacting populations and the eigenvalues are precisely the Lyapunov exponents that will govern the convergence to the possible equilibrium conditions. Then, near the equilibrium point (fixed point), we have

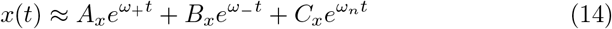

where *x* represents either *E*,*R* or *S*. At a stable equilibrium, all eigenvalues have negative real parts. The asymptotic long-term behavior of the system is governed by the Lyapunov exponent with the smaller real part (in modulus), say *λ*, i.e. *x*(*t*) *− x* ∝ e^λt^*.

Fig. 7a shows the time evolution of all species for *n* = 1 in the case of refuge mechanism. In the inset we show the long-term exponential convergence to the equilibrium populations. All of them are governed by a single Lyapunov exponent *λ* = *ω_n_*. Since the long-term rate of convergence to the coexistence fixed point is determined by a pure real eigenvalue, the trajectories shown in inset figure are purely exponential. In that case, similar results are recovered by any *n* within (0.0*, n\**]. For morphological defense, the other eigenvalues can take into place in the asymptotic regime. In Fig. 7b we display the same as Fig. 7a for morphology-induced mechanism with *n* = 8. The inset figure shows that the Lyapunov exponent is complex where the long-term behavior has oscillations in the exponential convergence. The precise value of the lowest Lyapunov exponent was determined from the Jacobian matrix for the internal and boundary fixed points. We calculated the Lyapunov exponents for various values of *n*, as shown in Fig. 8.

**Figure 7:**
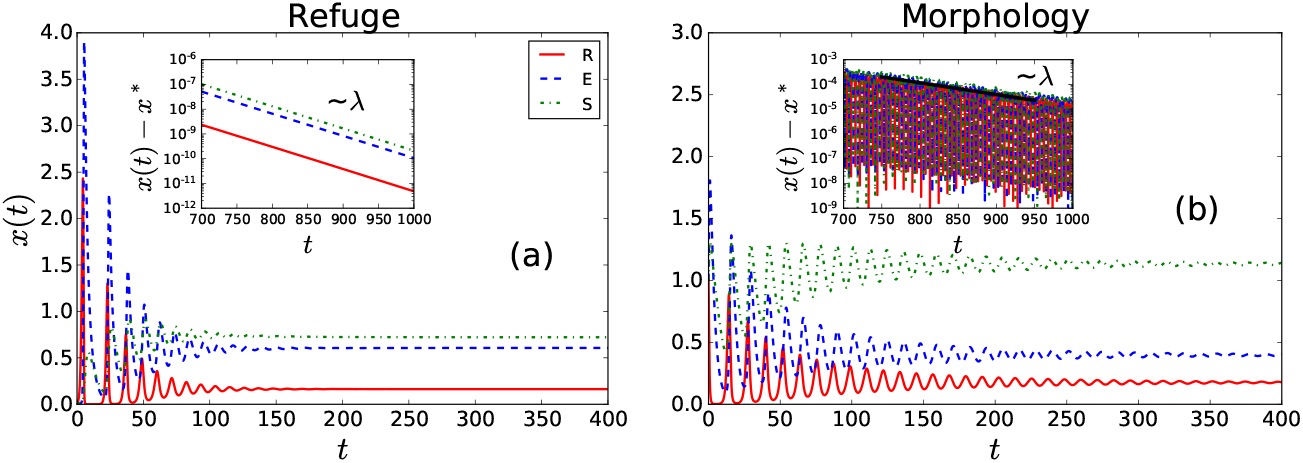
Time evolution of the population densities. (a) Refuge defense with *n* = 1. (b) Morphological defense with *n* = 8. The insets show the long-time behavior dominated by the lowest Lyapunov exponent *λ* (real for refuge and complex for morphological defense). Here *x* represents either *R*,*E* or *S*. Parameters are given in Table 1.

**Figure 8:**
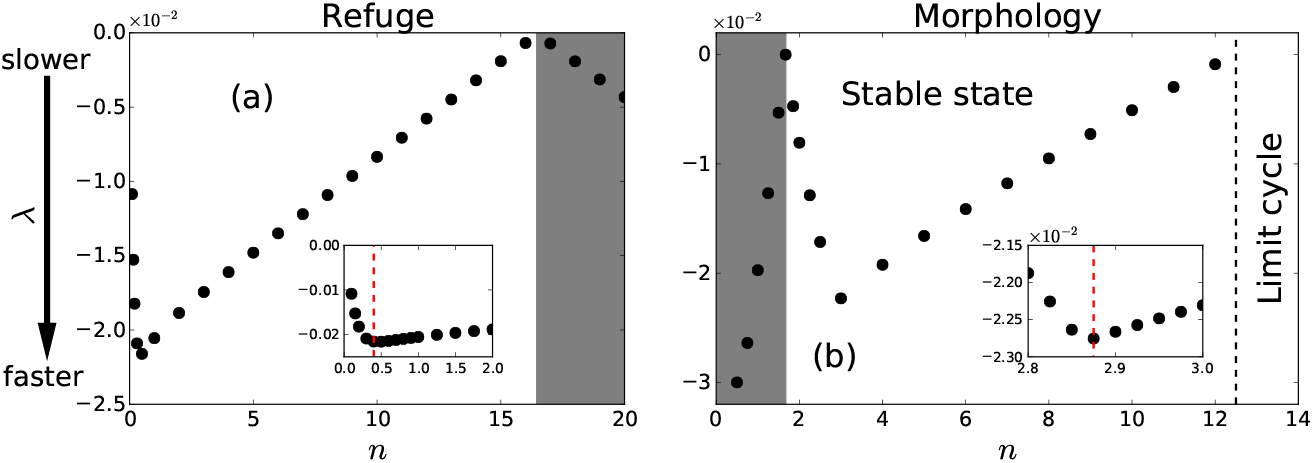
Lyapunov exponent against the degree of nonlinearity. Grey region indicates the deterioration of the food web characterized by the extinction of one IG population. (a) Refuge defense: Deterioration occurs above a characteristic value *n\** where *S* is extinct. The inset shows that the optimal Lyapunov exponent leading to the fastest convergence to coexistence occurs at *n* ≃ 0.45 (red dashed line). (b) Morphological defense: Below the characteristic value *n\** the IG-prey is extinct. The other phase is subdivided into two regions where the three-species community coexist in a stable fixed point or a limit cycle. The inset shows that the optimal Lyapunov exponent leading to the fastest convergence to coexistence occurs at *n ≃* 2.875 (red dashed line). Parameters are given in Table 1.

Two clear regimes arise from refuge model. With a strong nonlinearity degree *n > n\** (grey region) the top predator attack is so inefficient that the IG-prey population overcomes and *S* is extinct due to a shortage of basic resources. In this regime the Lyapunov exponent actually governs the rate of convergence towards the boundary (*R*, E*,* 0) fixed point. Below *n\** the Lyapunov exponent characterizes the rate of convergence towards the coexistence fixed point. It vanishes at *n\** as well as at *n* = 0 where the boundary fixed point (*R*,* 0*, S\**) becomes stable. Therefore, the rate of convergence to the coexistence equilibrium point depends non monotonically of the nonlinearity degree of the defense mechanism. There is a regime of moderate nonlinearities for which the rate of convergence grows with *n*. On the other hand, it becomes slower as *n* approaches *n\**. The threshold from weak to moderate nonlinearities can be delimited by the condition leading to the fastest convergence to the internal fixed point. For the parameters set used, it occurs roughly at *n* = 0.45.

The morphology-induced defense mechanism brings a richer range of dynamical regimes. Fig. 8b compile all of these different behaviors. As it was calculated analytically, at *n\** a stability switch occurs with the Lyapunov exponent vanishing at this point. The characteristic nonlinearity degree separates the phases of coexistence and of IG-prey population extinction. Right after *n\**, the asymptotic behavior is governed by the pure real Lyapunov exponent. Above a specific nonlinearity, the convergence becomes governed by the complex-valued Lyapunov exponents as in Fig. 7b. For the parameters set used, the fastest convergence to the internal fixed point occurs roughly at *n* = 2.875 as shown in the inset figure of Fig. 8b. Above an even larger value of *n*, the Lyapunov exponent of the internal fixed point becomes positive leading to an unstable fixed point. In this regime, the species coexist in a limit cycle oscillatory behavior (recall Fig. 4d).

### 3.3. Saturation effect in the refuge response

Due to the limited availability of refuge places, its consequent nonlinear influence on the attack efficiency shall saturate for large populations of IG-prey. Choosing a saturating function in order to model the effective refuge defense of the prey population, we are assuming the efficiency of this defensive form to be bounded. A proper saturable refuge defense can be modelled by considering the attack efficiency to have the form

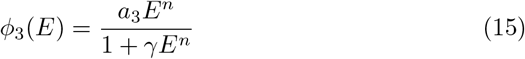

As seen in Fig. 9, changing *n* leads to a switch from Type II (*n* = 1) to Type III (*n >* 1) functional responses. As theorized by Huang et al. [36], even in the case of refuge mechanism, different species are represented by different functional responses. With extremely high *E* values, the response function saturates at its plateau *a*_3_*/γ*. Increasing the nonlinearity degree, the attack function goes to zero faster. In the same way, this increment tends to a faster saturation towards the maximum attack efficiency. As a consequence, we notice a possible trade-off in the degree of nonlinearity *n* and the saturation strength *γ*. The non-saturating model and its results are directly recovered with *γ* = 0. Furthermore, it is also recovered in the regime of small *E* (seen by a Taylor series expansion of Eq. 15). Then, the prior non-saturating refuge defense model is appropriated for systems where the IG-prey cannot grow large enough. The effect of saturation in the previous results is that it suppresses the regime of species coexistence. The characteristic value of the nonlinear strength *n\** delimiting the coexistence phase for this saturable refuge defense can be written as

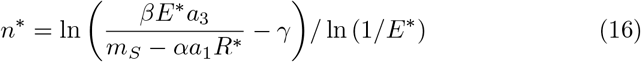

**Figure 9:**
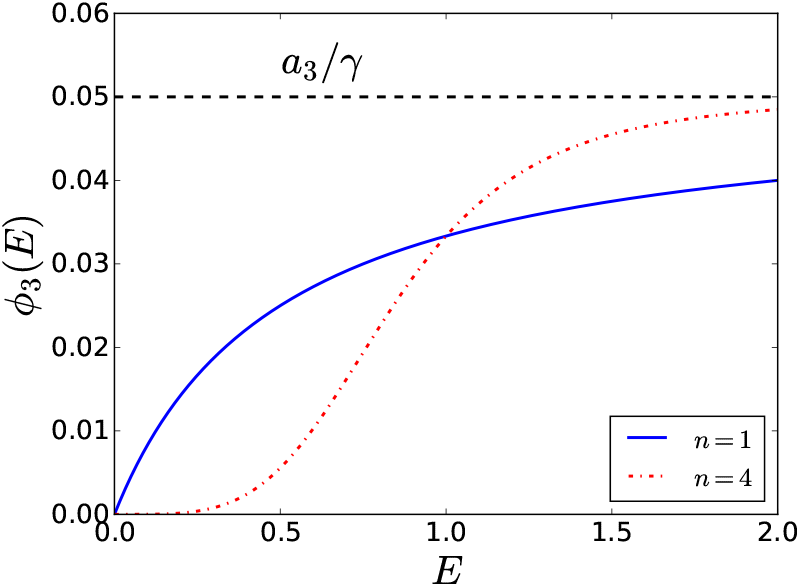
Saturable refuge response function as a function of IG-prey abundance for different nonlinearity degrees. It is clear that the Type II and Type III responses are recovered by *n* = 1 and *n >* 1, respectively. For this plot we set *a*_3_ = 0.1 and *γ* = 2.

where *R\** and *E\** are the same as Eq. 12. Fig. 10 shows the phase diagram for the saturating case where it is clearly unveiled that the range of coexistence decreases when *γ* is increased.

**Figure 10:**
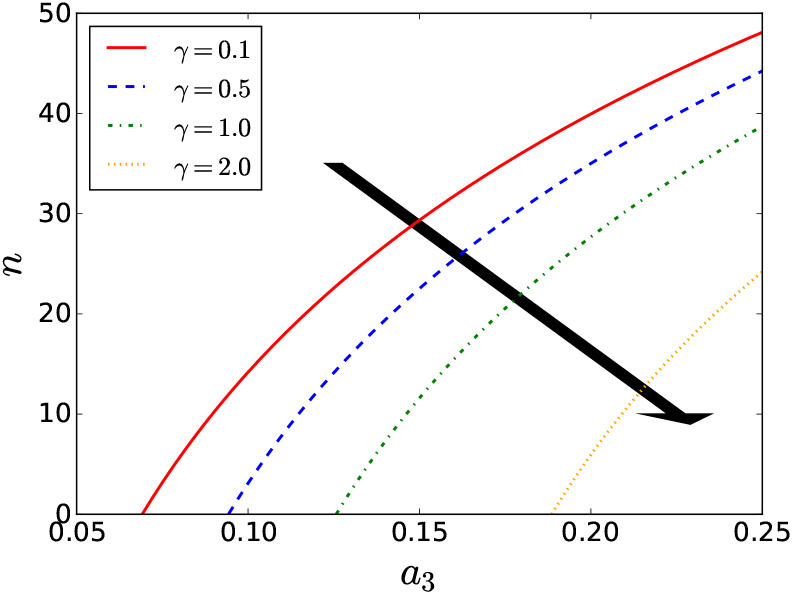
Phase diagram in the parameter space spanned by the attack efficiency *a*_3_ and nonlinearity *n* for a saturable refuge defense. Each line shows the characteristic nonlinearity *n\** which delimits the coexistence phase. Species can coexist below each curve. The range of values that allows coexistence decreases when increasing *γ*. Parameters are the same given in Table 1.

## 4. Summary and Conclusions

In summary, we introduced a three species population dynamics model in which a top predator and an intermediate species share the same resource. The dynamic equations only on rare occasions support a stationary coexistence solution of all three populations when the attack of the top predator to the intermediate species is linear. Motivated by the ubiquitous existence of coexistence food webs in nature and of experimental data reporting defense mechanisms of the intraguild species, we considered nonlinear attack terms to model either prey refuge or morphological defensive responses. As the degree of nonlinear defense *n* increases, the attack becomes more inefficient. For a refuge-like defense, the attack is compromised at low populations of the intraguild species. In the case of a morphological defense, it is triggered by a large predator population. Both defensive reactions favor a coexistence regime.

We provided a detailed analysis of the proposed models of a nonlinear interaction between the top predator and the intermediate prey. We demonstrated that the nonlinearity induced by the defense mechanism actually induces the emergence of a wide regime of species coexistence. For a refuge defense, the attack becomes so inefficient for very strong nonlinearities that it can lead to the extinction of the top predator population, assuming the IG-predator could not utilize prey-switching or adaptive feeding and that the remaining available basic resource is not enough to sustain it. Coexistence is found to be stable for 0 *< n < n\**, where *n\** was shown to grow logarithmically with the linear attack efficiency *a*_3_. We further unveiled that there is an optimal nonlinearity degree for which the rate of convergence towards the coexistence equilibrium is the fastest. This finding can be of evolutionary relevance since natural selection usually drives the interaction between surviving populations to the most adapted condition [32, 45, 46]. Saturation of the nonlinear reaction was shown to reduce its capability to induce species coexistence.

In the case of a morphological defense triggered by a large population of predators, there is a minimal nonlinearity required to induce the species coexistence. Very large nonlinearities can make the system dynamics evolves towards a limit cycle state with populations presenting persistent oscillations. Convergence to the stationary coexistence state is also faster at a finite nonlinearity. The models introduced here can be extended to more complex food web networks. It would be interesting to explore the role played by such nonlinear defense mechanism in other population dynamics as well as Webworld models [40, 41]. We hope the present work also motivate experimental works aiming to estimate the degree of nonlinear defense in intraguild predation food webs.

## Acknowledgements

This work was partially supported by CNPq, CAPES, and FINEP (Federal Brazilian Agencies), and FAPEAL (Alagoas State Agency). JPM acknowledges the hospitality of ICTP-SAIFR at IFT-UNESP during the *VII Southern Summer School on Mathematical Biology* where this project started.

